# NYUS.2: an Automated Machine Learning Prediction Model for the Large-scale Real-time Simulation of Grapevine Freezing Tolerance in North America

**DOI:** 10.1101/2023.08.21.553868

**Authors:** Hongrui Wang, Gaurav D. Moghe, Al P. Kovaleski, Markus Keller, Timothy E. Martinson, A. Harrison Wright, Jeffrey L. Franklin, Andréanne Hébert-Haché, Caroline Provost, Michael Reinke, Amaya Atucha, Michael G. North, Pierre Helwi, Michela Centinari, Jason P. Londo

**Author notes:** Author for correspondence Hongrui Wang (**) and Jason P. Londo (*).

## Abstract

- Accurate and real-time monitoring of grapevine freezing tolerance is crucial for the sustainability of the grape industry in cool climate viticultural regions. However, on-site data is limited. Current prediction models underperform under diverse climate conditions, which limits the large-scale deployment of these methods.
- We combined grapevine freezing tolerance data from multiple regions in North America and generated a predictive model based on hourly temperature-derived features and cultivar features using AutoGluon, an automatic machine learning engine. Feature importance was quantified by AutoGluon and SHAP value. The final model was evaluated and compared with previous models for its performance under different climate conditions.
- The final model achieved an overall 1.36 °C root-mean-square error during model testing and outperformed two previous models using three test cultivars at all testing regions. Two feature importance quantification methods identified five shared essential features. Detailed analysis of the features indicates that the model might have adequately extracted some biological mechanisms during training.
- The final model, named NYUS.2, was deployed along with two previous models as an R shiny-based application in the 2022-2023 dormancy season, enabling large-scale and real-time simulation of grapevine freezing tolerance in North America for the first time.

## 1. Introduction

The global and regional distribution of perennial plants is primarily constrained by abiotic stresses associated with regional climate. For example, the cultivation of European grapevines (*Vitis vinifera*) in mid-winter cold regions in North America presents a significant challenge, as the minimum temperatures in these regions sometimes exceed the plants’ maximum freezing tolerance. Cold-related damage is therefore a major limiting factor for the grape and wine industries in these regions (Zabadal *et al*., 2007; Poling, 2008; Dami *et al*., 2016; Londo & Kovaleski, 2017). Along with the development of preventative cultural practices and the introduction of cultivars with improved freezing tolerance, the monitoring of grapevine freezing tolerance has been a major focus of research groups in cool climate viticultural regions in North America to support the sustainability of the grape and wine industries.

Currently, the monitoring programs for grapevine freezing tolerance mainly rely on measuring the dormant bud low temperature exotherm (LTE), a burst of heat released when intracellular ice formation occurs, using a method called differential thermal analysis (DTA) (Pierquet & Stushnoff, 1980). DTA is conducted in programmable freezers, with buds exposed to a gradual decrease in temperature from 0 °C to a lethal temperature (−40 °C to −50 °C) at specific rates (e.g., −4 °C·h^-1^), and LTEs are recorded as a voltage change by a thermoelectric module placed under sample plates (Mills *et al*., 2006; Londo *et al*., 2023). Although this method facilitates a rapid assessment of bud freezing tolerance compared to a visual assessment of oxidative browning (Wample *et al*., 1990; Mills *et al*., 2006; Londo & Kovaleski, 2017), the whole procedure, along with sample collection and preparation typically takes days, which eliminates its large-scale application with high temporal resolution. As a result, most grapevine freezing tolerance monitoring systems established in cool climate viticultural regions in North America only report weekly or bi-weekly updated freezing tolerance of a few cultivars collected from a few sites. Although the accuracy is ensured, current DTA-based monitoring systems might not properly oblige the growers when climate change-associated weather extremes occur more frequently (Cohen *et al*., 2021). An advanced grapevine freezing tolerance prediction system that accommodates multi-site and multi-cultivar freezing tolerance with high temporal resolution could serve not only as a practical tool for vineyard management but also as a guideline for the adaptation of viticulture in the era of climate change.

To this end, there has been an increased interest in developing prediction models to mathematically estimate grapevine freezing tolerance under field conditions. The most widely used models, WAUS.1 and its successor WAUS.2 are discrete-dynamic models, where daily changes in the lethal temperature for 50% of a bud population (bud LT_50_) are phased with incremental time steps (Ferguson *et al*., 2011, 2014). These models estimate daily freezing tolerance change based on daily maximum and minimum temperatures and cultivar-specific parameters (Ferguson *et al*., 2011, 2014). A derivative of WAUS.2 was also developed for cold climate interspecific cultivars in Wisconsin using local freezing tolerance data (North *et al*., 2021). A new model, NYUS.1, is a biological model recently developed using phased integration of cold acclimation and deacclimation responses based on recent findings of dormancy-related grapevine freezing tolerance dynamics (Kovaleski *et al*., 2018, 2023; North *et al*., 2022). However, two issues compromise the widespread application of these models. First, the parameters used in the WAUS.2 and NYUS.1 models were optimized using local freezing tolerance data in Washington and New York, respectively, which may contribute to overfitting under local climate conditions and underperformance under other climate conditions. The underperformance of WAUS.2 has been observed in New York, Wisconsin, and in subtropical viticultural regions in Chile (Rubio & Pérez, 2020; North *et al*., 2021; Kovaleski *et al*., 2023). Second, since these models are the mathematical realizations of the biological understanding of dormancy and freezing tolerance, the parameters in the models such as the dynamics of cold acclimation and deacclimation need to be updated regularly according to the advances in dormancy and freezing tolerance biology. The version update of these models can involve substantial manual parameter generation, selection, and tuning, which in turn compromises the deployment of the most updated version.

Machine learning (ML) and its subfield deep learning (DL) have significantly advanced in the last decade (Zhou, 2021; Ashenden *et al*., 2021). ML algorithms are typically used for structured data, such as numerical data in tables, which usually involves manual feature extraction. DL algorithms are capable of processing unstructured data, such as images, audio, text and sequence, which often require higher-level abstractions and more training data to be fully functional (Janiesch *et al*., 2021). Regarding the modeling of weather-related plant physiological responses, large scale applications of ML and DL are used for growth and yield estimation of major crops, such as corn, wheat and soybean (Khaki & Wang, 2019; Khaki *et al*., 2020; Shahhosseini *et al*., 2020; Gall *et al*., 2022; Ma *et al*., 2022). Researchers recently used a DL tool, RNN (recurrent neural network), as a backbone to model grapevine freezing tolerance. Being trained with Washington state data, the RNN model performed equivalently or better for some cultivars than WAUS.2 (Saxena *et al*., 2022). The model has yet to be evaluated in other regions due to ongoing further development. Nevertheless, the modeling of plant physiology using ML is still at very early stages. One leading reason is that the rapid development of ML can make it difficult even for ML experts to efficiently incorporate novel practices and timely deployed ML models. Another reason is that the training datasets are usually small due to the complex and time-consuming methods to measure plant physiological responses, such as measuring freezing tolerance with DTA (Feng *et al*., 2020; Yoosefzadeh-Najafabadi *et al*., 2021). The individual prediction models generated from such small datasets are subjected to overfitting, which limits their applications in real-world scenarios (Ali *et al*., 2014). The development of automated machine learning (Auto-ML) frameworks addresses some of these issues. These frameworks not only fuse all the tedious steps of ML model development, including data preprocessing, feature engineering, model training and hyperparameter tuning into very few lines of code, but also incorporate novel techniques such as model stacking and weighting to minimize overfitting. These tools enable domain experts, such as plant physiologists, to build ML applications through an automated pipeline without much requirement for coding and ML knowledge (He *et al*., 2021).

In this study, we combined the on-site measurement of freezing tolerance data (LT_50_) from multiple regions in North America and labeled each datapoint with 117 features generated from hourly temperatures and cultivar information. The dataset was used for the training of a prediction model through an Auto-ML framework, AutoGluon (Erickson *et al*., 2020). AutoGluon is hosted by Amazon Web Services (AWS) and is known for its ease of use and high performance through built-in multi-layer stacking and weighting techniques (Fakoor *et al*., 2020; Erickson *et al*., 2020). The first objective of this study was to build a generalizable prediction model that accurately estimates the grapevine freezing tolerance under different climate conditions. The second objective was to quantify the importance of each feature in the final model to understand the physiological control of freezing tolerance in grapevine. The third objective was to apply the model to cool climate viticultural regions to achieve a large-scale and real-time monitoring of grapevine freezing tolerance with high temporal resolution.

## 2. Materials and Methods

### 2.1 Grapevine freezing tolerance data and weather data

On-site measurements of grapevine bud freezing tolerance from multiple viticultural regions in North America were used for model training and evaluation. In all data collection regions, the freezing tolerance of a cultivar on a single date was determined using multiple buds through standard DTA with slight variations between collaboration groups (Mills *et al*., 2006; Londo *et al*., 2023). Briefly, grapevine buds were excised from canes collected from the field at specified time throughout the dormancy season and subjected to a decrease of temperature from 0 °C to a lethal temperature (−40 °C to −50 °C) in programmable freezers, and thermoelectric modules detects LTE that corresponds to the freezing tolerance of the buds. Bud LT_50_ was determined by either taking the mean or median of all LTEs, or through probit analysis. Slight variations in protocol do not significantly affect measurements in grapevine (Londo *et al*., 2023). The LT_50_ dataset composition is summarized in Fig. 1. The dataset combines the LT_50_ measurements from the nine grapevine freezing tolerance programs in New York (NY), Michigan (MI), Wisconsin (WI), Washington (WA), Pennsylvania (PA) and Texas (TX) in the U.S. and British Columbia (BC), Québec (QC) and Nova Scotia (NS) in Canada between 2002 to 2023 (Fig. 1a). These regions cover the major types of climate conditions in the cool climate viticultural regions in North America. The dataset contains 3,458 LT_50_ measurements from 24 *Vitis* sp. hybrid cultivars and 6,699 LT_50_ measurements from 21 *Vitis vinifera* cultivars, resulting in a total of 10,157 datapoints (Fig. 1b and c). The 45 cultivars in this dataset represent a large proportion of the cultivars grown in North America. Hourly or daily temperature data were obtained from various sources (Table S1) to represent local weather conditions at the sites of field data sampling. At the sites where hourly temperature was unavailable, hourly temperature was estimated based on daily maximum and minimum temperature using ‘*stack_hourly_temps*’ of the R package ‘chillR’ (Luedeling & Fernandez, 2022).

**Fig. 1.**
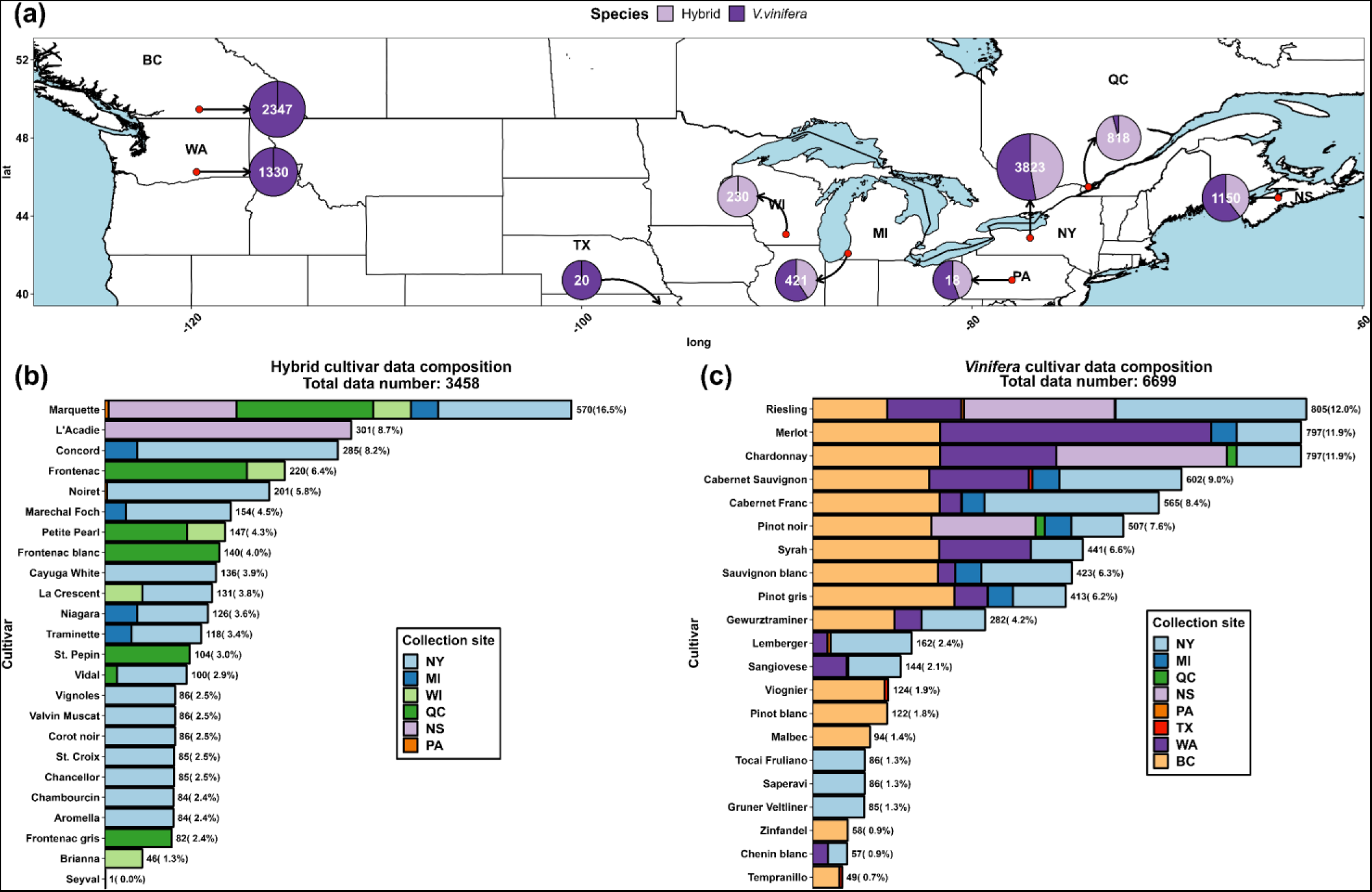
Grapevine freezing tolerance (LT50) dataset composition. (a) Geographic distribution of data collection sites. NY dataset was obtained from Cornell University Cornell AgriTech, Geneva, NY, USA (42.88° N, 77.03° W) and composed of 3,823 LT50 data. Among these data, 1,807 (47%) were collected from hybrid cultivars, and 2,016 (53%) were collected from *V. vinifera* cultivars. MI dataset was obtained from Michigan State University Southwest Michigan Research and Extension Center, Benton Harbor, MI, USA (42.08° N, 86.36° W) and composed of 435 LT50 data. Among these data, 183 (42%) were collected from hybrid cultivars, and 252 (58%) were collected from *V. vinifera* cultivars. WI dataset was obtained from University of Wisconsin Madison, West Madison Agricultural Research Station, Verona, Wisconsin, USA (43.06° N, 89.53° W) and composed of 230 LT50 data collected from hybrid cultivars. WA dataset was obtained from Washington State University, Irrigated Agriculture Research and Extension Center, Prosser, WA, USA (46.25° N, 119.74° W) and composed of 1,330 data collected from *V. vinifera* cultivars. PA and TX datasets are two minor datasets contributed by The Pennsylvania State University and Texas A&M University AgriLife Extension Service, respectively. BC dataset was obtained from Agriculture and Agri-Food Canada (AAFC), Summerland Research and Development Center, Summerland, BC, Canada (49.46° N, 119.60° W) and composed of 2,347 LT50 data collected between 2012 to 2022 from 15 *V. vinifera* cultivars. QC dataset was obtained from Centre de Recherche Agroalimentaire de Mirabel, Mirabel, QC, Canada (45.49° N, 74.05° W) and composed of 818 LT50 data. Among these data, 784 (95%) were collected from hybrid cultivars, and 34 (5%) were collected from *V. vinifera* cultivars. NS dataset was obtained from AAFC, Kentville Research and Development Centre, Kentville, NS, Canada (44.93° N, 64.32° W) and composed of 1,150 LT50 data. Among these data, 457 (40%) were collected from hybrid cultivars, and 693 (60%) were collected from *V. vinifera* cultivars. (b) Data composition of hybrid cultivars. (c) Data composition of *V. vinifera* cultivars.

### 2.2 Feature generation and extraction

The full model dataset contains 117 features. The features used for modeling in this study are within two categories: cultivar features and hourly temperature-based features, except for days in season, which is the number of days after September 1^st^. Cultivar features were generated by transforming single-column categorical cultivar names to multiple-column binary cultivar names through one-hot encoding (Ashenden *et al*., 2021). This approach addresses the difficulties of managing mixed-type variables in machine learning and facilitates the application of model interpretation techniques (Janiesch *et al*., 2021). Forty-five Boolean-type features were obtained from this transformation. The remaining 72 features are temperature and time related.

Hourly temperature-based features comprise daily temperature descriptors, cumulative temperature descriptors, exponential weighted moving average (EWMA) temperatures and reverse exponential weighted moving average (REWMA) temperatures. Daily temperature descriptors and cumulative temperature descriptors are detailed in Note S1. EWMA and REWMA temperatures are unique temperature descriptors computed based on temperature windows (Holt, 2004). The mathematical expression of EWMA and REWMA, along with their impacts on daily mean, maximum and minimum temperatures, are described in Fig. 2. EWMA is a smoothing method used for normalization during data preprocessing and to decrease variation in time series data in ML (Janiesch *et al*., 2021). In this study, the EWMA temperature of a date 𝑡 is computed based on a temperature window of a size 𝑛 (in days) with exponentially ascending weight added to each date from 𝑡 − 𝑛 to 𝑡 (Fig. 2a). This method allows all the temperatures in the window to contribute to the final EWMA temperature, while the closer the date is to date 𝑡, the more impact the temperature of the date has on the final EWMA temperature. As the window size 𝑛 increases, the EWMA temperatures follow the same trend as the original temperatures, but they are increasingly smoothed as more data are included in the weighted average (Fig. 2b).

**Fig. 2.**
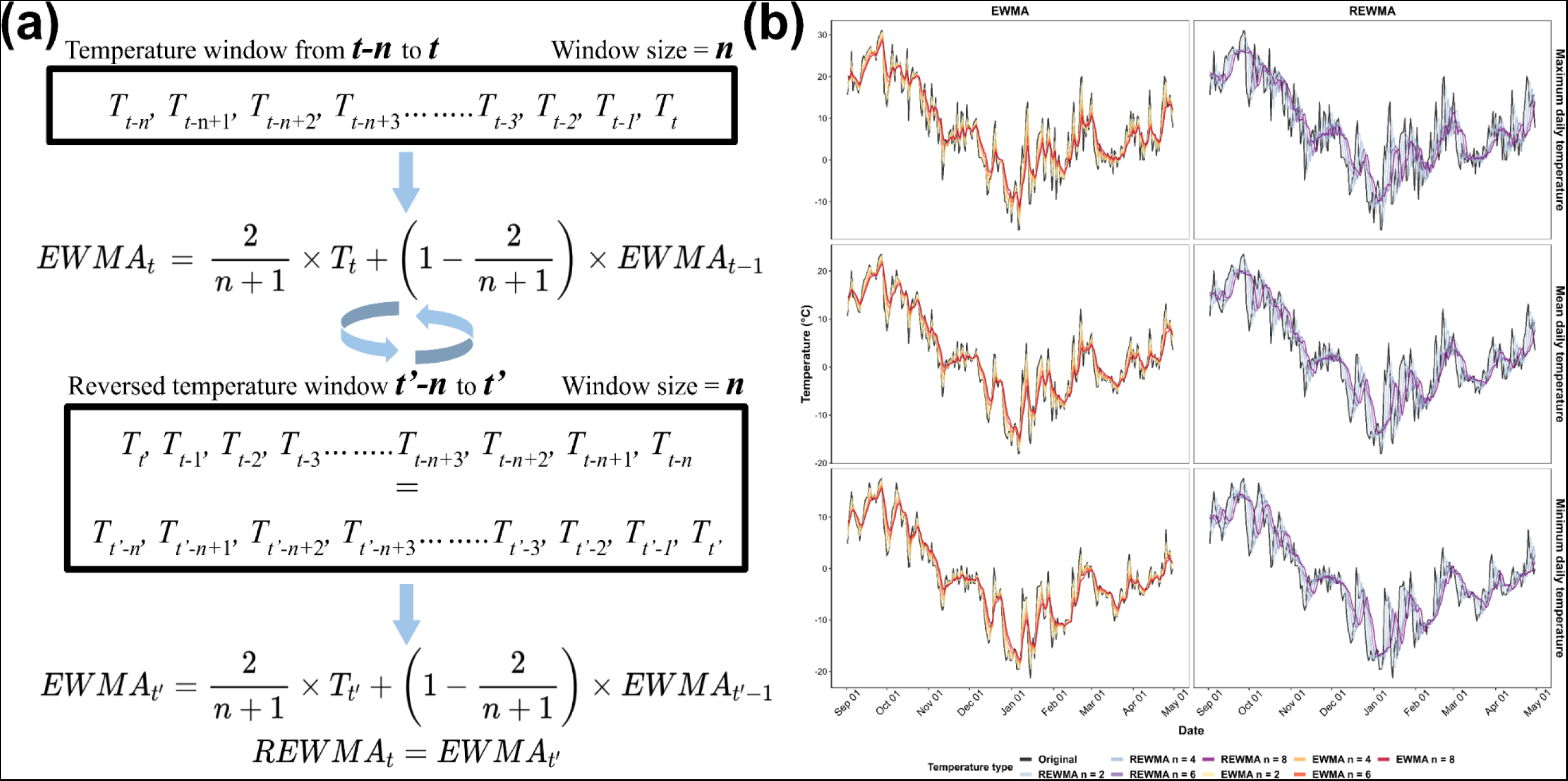
Computation of exponential weighted moving average (EWMA) and reverse exponential weighted moving average (REWMA) temperatures and their impacts on daily temperatures. (a) Schematic representation of EWMA and REWMA temperatures in a window from 𝑡 − 𝑛 to 𝑡. 𝑇 is the temperature of a day. To compute the EWMA of a date 𝑡 with the window size of 𝑛, weight is exponentially ascendingly added from 𝑡 − 𝑛 to 𝑡. To compute the REWMA of a date 𝑡 with the window size of 𝑛, the window is first reversed to 𝑡′ − 𝑛 to 𝑡′ (where 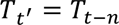 and 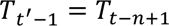, etc.), and weight is exponentially ascendingly added from 𝑡′ − 𝑛 to 𝑡′, thus exponentially descendingly added from 𝑡 − 𝑛 to 𝑡. (b) Effect of EWMA and REWMA with four window sizes (2, 4, 6 and 8 days) on maximum, mean and minimum daily temperatures in a dormant season. Original temperatures represent the recorded temperature in the field during the season.

Including EWMA temperatures as features allows Auto-ML to factor temperature as a cumulative descriptor from a temperature window rather than only relying on daily temperatures. This approach may result in higher accuracy since the change of freezing tolerance is more likely a dynamic consequence of continuous exposure to a temperature window rather than a transient response to current temperature (De Rosa *et al*., 2021).

REWMA temperature is a new expression of temperature to incorporate the theory of ‘cold priming’ and the ‘cold shock effect’ in the modeling of grapevine freezing tolerance. The biological reasoning for including REWMA temperatures is available in Note S2. The major difference between REWMA and EWMA is that the assignment of weight is reversed in REWMA, thus allowing earlier temperature to have more impact. To compute REWMA, the temperature window (size = 𝑛) from date 𝑡 − 𝑛 to date 𝑡 (𝑇_𝑡−𝑛_ to 𝑇_𝑡_, where 𝑇 is the temperature of a day) is reversed. EWMA is applied on the reversed window (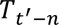 to 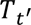, where 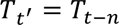 and 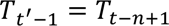, etc.) (Fig. 2a). In this method, weight exponentially descends from 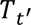 to 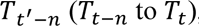, whereas all the temperatures in the window contribute to the final 𝑅𝐸𝑊𝑀𝐴_𝑡_. Similar to EWMA, as window size 𝑛 increases, the REWMA temperatures are smoothed as more data is used for computation. However, in REWMA, all the temperatures also better correlate with earlier dates (visually horizontally shifted) as compared to the original temperatures (Fig. 2b).

We extracted EWMA and REWMA temperatures of daily maximum, minimum and mean temperatures with window sizes in two-day increments (2:20) for a total of 60 features. A correlation matrix of the EWMA and REWMA of daily minimum temperature is shown in Fig. S1. As the window size increases, the correlations between all these temperature features decrease (Fig. S1). The correlations between REWMA temperatures are generally weaker than those between EWMA temperatures, and the correlations between EWMA temperatures and REWMA temperatures are even weaker (Fig. S1).

### 2.3 Method suitability test, site-transferability test, model training and model selection

Since this study represents the first approach to grapevine freezing tolerance modeling using Auto-ML, the first step was to test for method suitability and site-transferability. We therefore performed three types of validation: (i) cross-validation during model training, (ii) validation using an internal testing dataset (‘internal’) and (iii) validation using an external testing dataset (‘external’). Specifically, the LT_50_ data from three minor sub-datasets in MI, PA and TX (𝑛 = 459) were isolated and used as external testing data. The remaining data (𝑛 = 9,698) were used to train and test an alpha grapevine LT_50_ prediction model for the method suitability test and the site-transferability test. The partial dataset (𝑛 = 9,698) was randomized before training to avoid structural bias and split into 90% and 10% subsets to be used for training (𝑛 = 8,728) and internal testing (𝑛 = 970), respectively. Only the training data was used for model training and ten-fold cross-validation. In this study, the difference between internal testing data and external testing data is that the internal testing data were from the same collection regions as the training data (thus named ‘internal’), and the external testing data were from the different collection regions (thus named ‘external’). The performance of the alpha model on the internal testing data informs if Auto-ML can generate a model that can accurately predict the LT_50_ of the data that were not used for training (𝑛 = 970), which indicates ‘method suitability’. In comparison, the performance of the alpha model on the external testing data informs whether the resulting model can accurately predict the LT_50_ of the data that were neither used for training nor originated from the same regions (𝑛 = 459), which indicates ‘site-transferability’.

The Python package AutoGluon (version 0.7.0) (Erickson *et al*., 2020) was used to conduct model training. Training was conducted through ‘*TabularPredictor*’ with ‘*fit(presets=’best_quality’, num_bag_folds = 10, num_stack_levels = 5)*’. Root-mean-square error (𝑅𝑀𝑆𝐸) was used as the evaluation metric for model performance. All steps of data preprocessing were automatically managed by AutoGluon. All the models produced from training were evaluated with 𝑅𝑀𝑆𝐸 using internal testing data. As the training command specifies stacking and weighting at five levels (*num_stack_levels = 5*), it is possible that model performance maximizes after certain stacking levels. In this case, the model with the least complexity (lowest level of stacking) was selected as the resulting model per training to enable highest prediction speed. After the method suitability test and the site-transferability test, the entire dataset was aggregated to include LT_50_ from every region (𝑛 = 10,157) to generate a final grapevine LT_50_ prediction model. The training of the final model was the same as stated above, but the dataset was randomized again to be split into 90% training data and cross-validation data (𝑛 = 9,141) and 10% internal testing data (𝑛 = 1,016).

### 2.4 Feature importance quantification of the final model

The importance of each feature in the final model was quantified using internal testing data through two methods. First, AutoGluon internal feature importance quantification was conducted using ‘*feature_importance*’ with ‘*(num_shuffle_sets=100, subsample_size=1000)*’. In this method, when testing data is perturbed by randomly shuffling the values of a feature across rows, the feature’s importance score reflects the resulting decrease in the model’s performance, a derivation of local interpretable model-agnostic explanations (LIME) (Ribeiro *et al*., 2016). Second, feature importance was quantified by computing the SHAP (SHapley Additive exPlanations) value of the feature using the Python package SHAP (version 0.41.0) (Lundberg & Lee, 2017). The SHAP value of a feature represents the average marginal contribution of that feature across all possible feature combinations. The calculation of SHAP values involves computing the contributions of each feature subset, considering all possible permutations of features (Štrumbelj & Kononenko, 2014).

### 2.5 Model comparison

To further test model performance, the final model was compared to two previous grapevine LT_50_ prediction models, WAUS.2 and NYUS.1. These models were generated using Washington LT_50_ data and New York LT50 data, respectively. (Ferguson *et al*., 2014; Kovaleski *et al*., 2023). Since the current version of NYUS.1 is limited to the LT50 of cultivars ‘Cabernet Sauvignon’, ‘Riesling’ and ‘Concord’ (Kovaleski *et al*., 2023), model performance comparison was conducted on a subset of internal testing data (𝑛 = 1,016) of ‘Cabernet Sauvignon’ (𝑛 = 69), ‘Riesling’ (𝑛 = 60) and ‘Concord’ (𝑛 = 31) from NY, WA, BC and MI sub-datasets.

### 2.6 Model deployment

In the 2022-2023 dormant season, the final model was applied on weather station temperature data and interpolated gridded temperature data in Applied Climate Information System (ACIS, https://www.rcc-acis.org/) to facilitate real-time grapevine freezing tolerance monitoring in cool climate viticultural regions in the U.S. The model prediction was updated daily and interactively deployed using the R Shiny package (Chang *et al*., 2022). The current monitoring system covers the 2,035 weather stations in Global Historical Climatology Network (GHCN, in ACIS) within the bounding box of Minnesota (NW), Maine (NE), Virginia (SE)and Missouri (SW). Gridded prediction is also available for Northeastern and Midwestern U.S. with a 5 km × 5 km spatial resolution based on interpolated gridded temperature data from ACIS. Along with LT50 prediction, the potential that freezing damage has occurred (0 to 100), is estimated through a symmetric sigmoid function.

The sigmoid function is defined as:

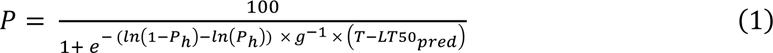

where, 𝑃 is the potential of freezing damage for a day, 𝑇 is the daily minimum temperature and 𝐿𝑇50_𝑝𝑟𝑒𝑑_ is the predicted LT_50_ from the model. 𝑃_ℎ_and 𝑔 are two constants that represent a potential of freezing damage that is greater than 50% and the gap of 𝑇 and 𝐿𝑇50_𝑝𝑟𝑒𝑑_ to reach that potential (Badr *et al*., 2018). In our estimation, we used 𝑃_ℎ_ = 0.9 and 𝑔 = 2, assuming that 10% and 90% of the potential would have occurred when the ambient temperature is 2 °C above and below the predicted LT_50_, respectively.

## 3. Result

### 3.1 Method suitability test and site-transferability test

Training results, model performances in internal testing and external testing for the method suitability test, and the site-transferability test are shown in Fig. S2. During training, five levels of stacking and weighting were accomplished, resulting in five weighted ensemble models and 47 stacker models at different levels (Fig. S2a). Internal testing indicated that multiple models (from ‘WeightedEnsemble_L2’ and the models thereafter) reveal the best performance with 𝑅𝑀𝑆𝐸 = 1.34 °C to 𝑅𝑀𝑆𝐸 = 1.37 °C (Fig. S2a). Thus, ‘WeightedEnsemble_L2’ (𝑅𝑀𝑆𝐸 = 1.35 °C) was selected as the alpha model for external testing for its least complexity. In the external testing, the alpha model achieved 𝑅𝑀𝑆𝐸 = 1.59 °C, 𝑅𝑀𝑆𝐸 = 1.44 °C and 𝑅𝑀𝑆𝐸 = 1.86 °C in the MI sub-dataset (𝑛 = 421), the PA sub-dataset (𝑛 = 18) and the TX sub-dataset (𝑛 = 20), respectively (Fig. S2b). The 𝑅𝑀𝑆𝐸 of individual cultivars with more than ten LT_50_ values ranges from 1.14 °C to 2.02 °C and is greater in several hybrid cultivars (e.g., ‘Concord’ and ‘Marquette’) (Fig. S2b).

### 3.2 Final model performance and feature importance

After these tests, the complete dataset (𝑛 = 10,157) was used to train and test the final grapevine LT_50_ prediction model. The complete dataset, along with date and site information, is provided in Data S1. Training data and internal testing data are provided in Data S2 and Data S3, respectively. Training results, model performance and feature importance quantification using AutoGluon and SHAP value are shown in Fig. 3. During training, multiple models (from ‘WeightedEnsemble_L2’ and the models thereafter) reveal the best performance (Fig. 3a), and WeightedEnsemble_L2’ (𝑅𝑀𝑆𝐸 = 1.36 °C) was selected as the final grapevine LT_50_ prediction model for its least complexity. The performance of the final model was visualized within each region (Fig. 3b). PA and TX data were omitted due to the limited amount of data categorized as internal testing data. The final model performance varies between 𝑅𝑀𝑆𝐸 = 0.82 °C (BC sub-dataset) and 𝑅𝑀𝑆𝐸 = 1.68 °C (NY sub-dataset (Fig. 3b). The final model is an ensemble model at stacking level two and is composed of eight base models generated at stacking level one. The prediction of the ensemble model is a weighted outcome of the predictions in the eight base models, and the weight of each base model is determined on their performance during training. The final model prediction on the LT_50_ of all internal testing data is provided in Data S4.

**Fig. 3.**
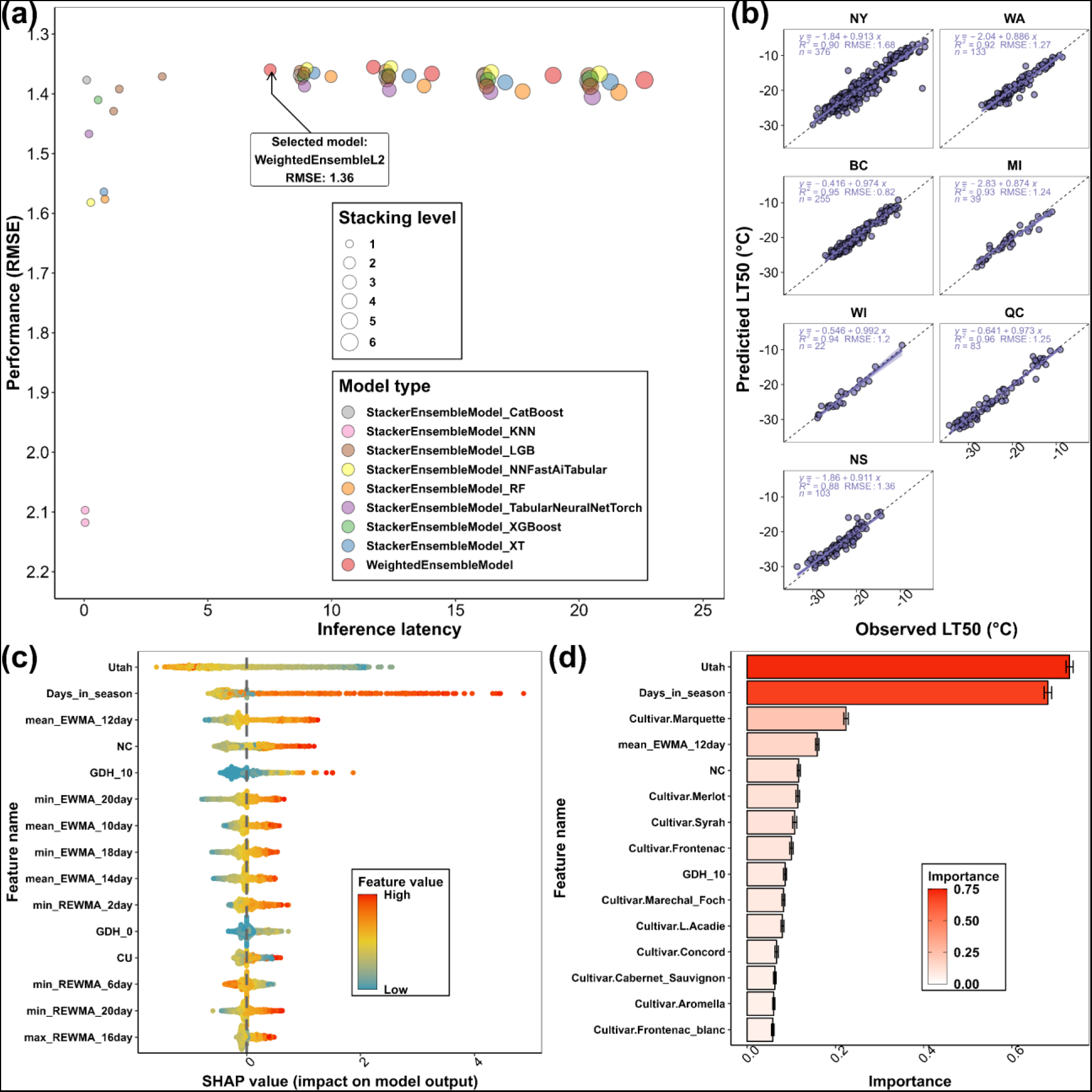
The final grapevine freezing tolerance model training result, performance, and feature importance. (a) The performance (RMSE) of all the models generated from training. Inference latency refers the relative time it takes for a machine learning model to process and analyze data and produce an output. (b) The performance of the final grapevine LT50 prediction model by individual sub-datasets. The data used to compute model performance are from internal testing data. (c) SHAP value distribution of the top 15 features of the final model. Feature importance is ranked based on the mean of absolute SHAP values of all the samples per feature. (d) AutoGluon feature importance of the top 15 features of the final model. The importance indicates the drop in model performance when the values of the feature are randomly shuffled across rows.

Feature importance was quantified using internal testing data (𝑛 = 1,016) through SHAP value and AutoGluon. The 15 most influential features identified in SHAP include three chilling models, two growing degree hours (GDH) models, nine EWMA/REWMA temperature features and days in season (Fig. 3c). The 15 most influential features identified by AutoGluon include two chilling models, one GDH model, one EWMA temperature, ten Boolean type cultivar features and days in season (Fig. 3d). Utah chilling hour model and days in season are the leading features that have far more impact on model performance (Fig. 3d). Randomizing the values of these two features across rows would result in 0.73 and 0.68 decrease of model performance (0.73 °C and 0.68 °C increase of 𝑅𝑀𝑆𝐸). Complete SHAP values of each internal testing data and each feature are provided in Data S5. A complete AutoGluon feature importance list is provided in Data S6.

### 3.3 Model performance comparison

The final model is thereafter referred to as ‘Auto-ML model’ in this study. The Auto-ML model was subjected to model performance comparison with NYUS.1 and WAUS.2. The results of the model performance comparison are shown in Fig. 4a. The prediction error is calculated by 𝐿𝑇50_𝑝𝑟𝑒𝑑𝑖𝑐𝑡𝑒𝑑_ − 𝐿𝑇50_𝑜𝑏𝑠𝑒𝑟𝑣𝑒𝑑_. As LT_50_ refers to subzero temperatures and thus is a negative value, negative prediction error indicates an overestimation, and positive prediction error indicates an underestimation of LT_50_. The shade of the prediction error distribution represents the 95% confidence interval of the trendline fitted using ‘loess’ and can serve as a rough measure of error-proneness. The temporal distribution of prediction error in the testing data is combined across all the dormant seasons and shown in Fig. 4b. The Auto-ML model prediction error is always relatively stable at around 0 °C in these three cultivars, but the confidence interval tends to be wider in late season (i.e., early spring), indicating relatively lower model stability as the buds approaching budbreak (Fig. 4b). NYUS.1 usually underestimates the freezing tolerance of ‘Cabernet Sauvignon’ and ‘Riesling’ but overestimates the freezing tolerance of ‘Concord’ (Fig. 4b). Significant prediction errors were identified in late season, and these errors primarily occurred in the predictions of WA, BC and MI data, suggesting a regional aspect that is being poorly predicted (Fig. 4b). WAUS.2 always overestimated the freezing tolerance of the three cultivars, and the overestimation became more severe in late season (Fig. 4b).

**Fig. 4.**
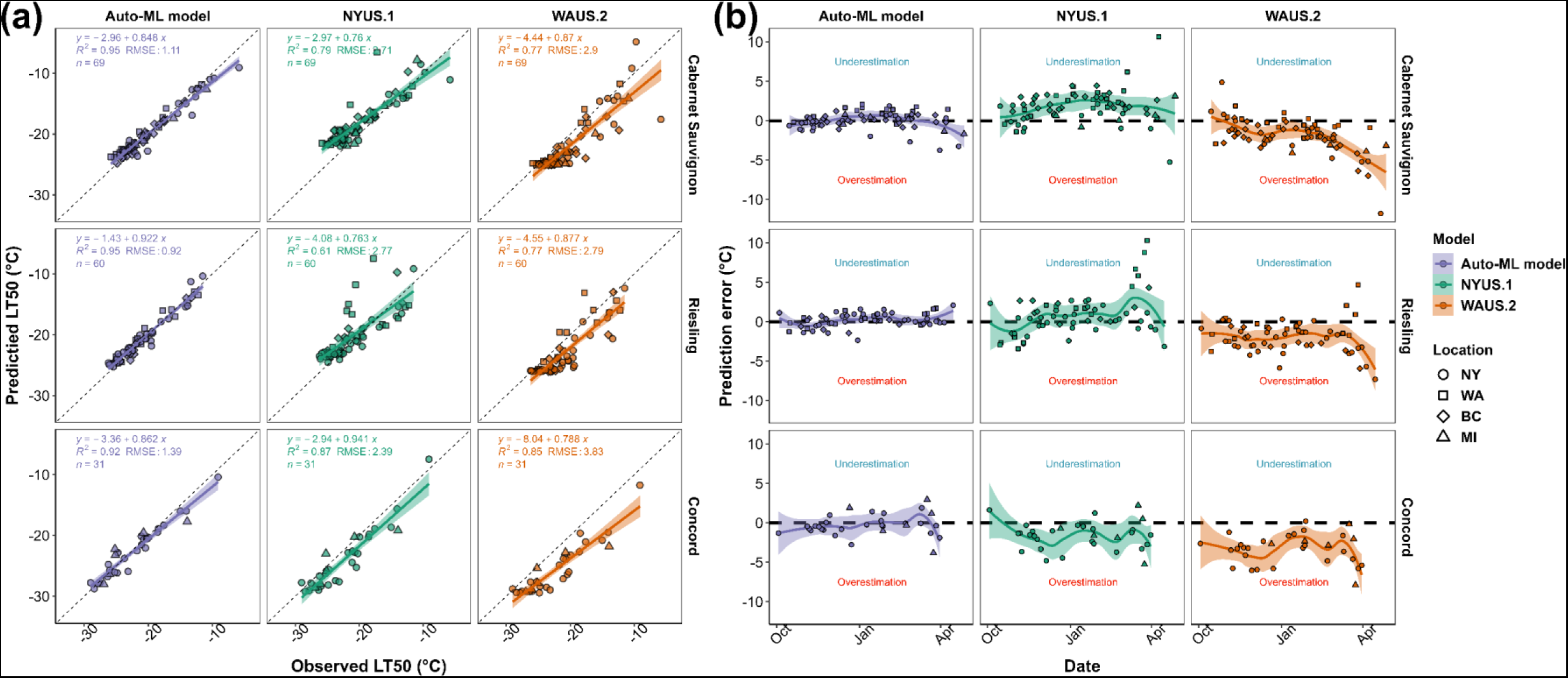
Grapevine freezing tolerance model comparison result and prediction error distribution. (a) The performance (𝑅𝑀𝑆𝐸) of Auto-ML model, NYUS.1 and WAUS.2 in ‘Cabernet Sauvignon’, ‘Riesling’ and ‘Concord’ in internal testing data. (b) Prediction error distribution by model and cultivar. Prediction error is calculated by 𝐿𝑇50_𝑝𝑟𝑒𝑑𝑖𝑐𝑡𝑒𝑑_ − 𝐿𝑇50_𝑜𝑏𝑠𝑒𝑟𝑣𝑒𝑑_. Negative prediction error indicates an overprediction, and positive prediction error indicates an underprediction. The confidence interval of the prediction error distribution can serve as a rough measure of error-proneness. These internal testing data were from NY (2012-2022), WA (2005-2012), BC (2012-2022), and MI (2021-2022) sub-datasets and were neither used for training nor cross-validation.

### 3.4 Model deployment

During the 2022-2023 dormant season, the Auto-ML model was deployed in the major cool climate viticultural regions in the U.S. through a new freezing tolerance website (https://grapecoldhardiness.shinyapps.io/grape_freezing_tolerance/). Predicted LT_50_ values throughout the dormancy season were made available for 16 cultivars at each weather station. Multiple sites in NY contributed on-site measurements of grapevine LT_50_ during the season through the data upload portal. Among these sites, Portland, NY (station ID: ‘USC00306747’) and Geneva, NY (station ID: ’USC00303184’) contributed whole season LT_50_ measurements. The predictions from Auto-ML model, NYUS.1 and WAUS.2 were compared to these measurements (Fig. 5). We note that these measurements were not used at all in the model development and were completely unseen by the model.

**Fig. 5.**
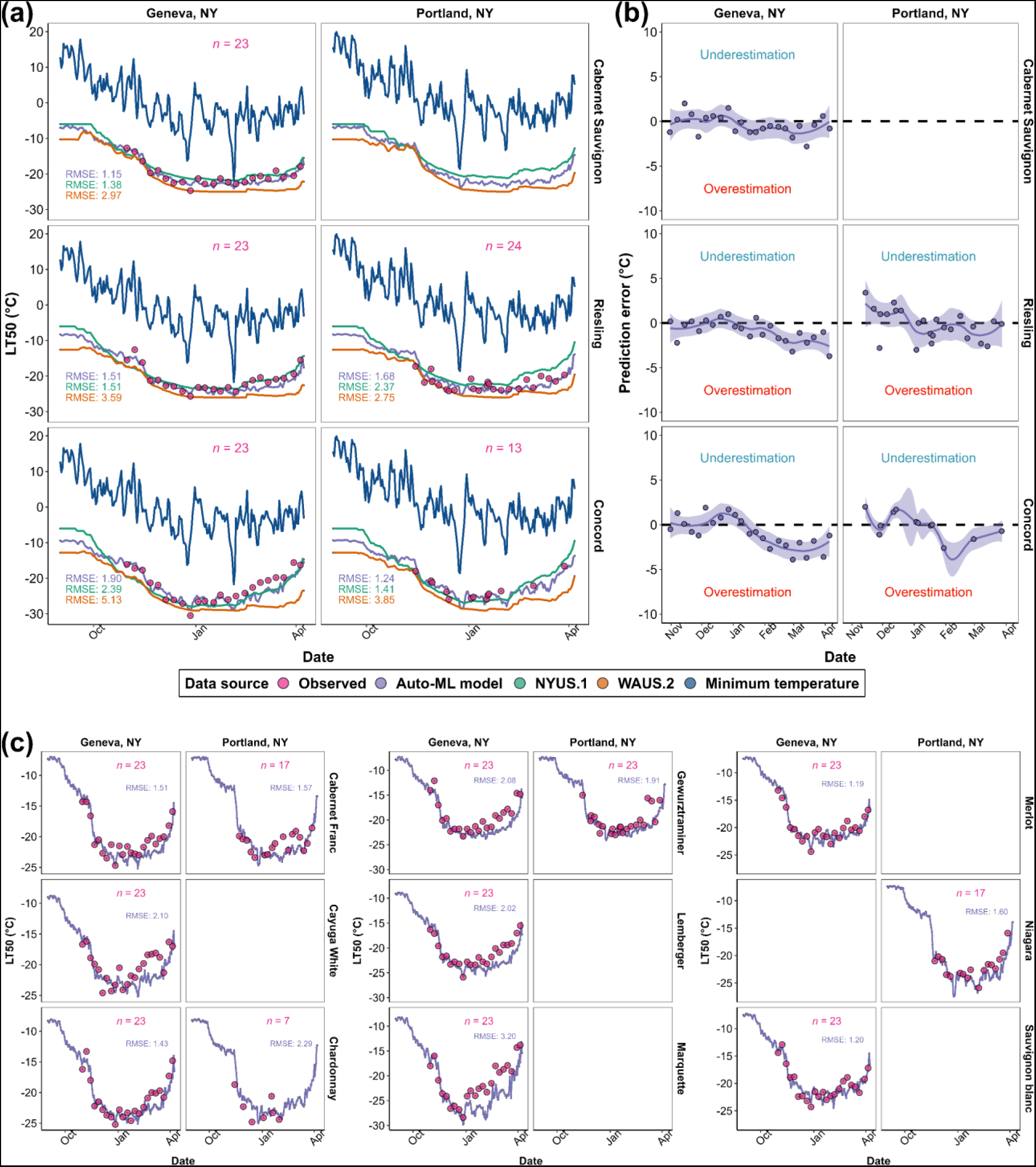
Model deployment result in Geneva, NY and Portland, NY in 2022-2023 dormant season. (a) The comparison of the predictions from Auto-ML model, NYUS.1 and WAUS.2 and on-site measurements in ‘Cabernet Sauvignon’, ‘Riesling’ and ‘Concord’. (b) Prediction error distribution of Auto-ML model by cultivar. (c) Predictions from Auto-ML model and on-site measurements in other cultivars.

During the season, the predicted LT_50_ values from the Auto-ML model prediction were typically intermediate to those predicted by WAUS.2 and NYUS.1 models and were more responsive to the change of minimum temperature (Fig. 5a). The Auto-ML model outperformed the previous models for all three cultivars at both sites (Fig. 5a). Prediction error distribution for the Auto-ML model across the season indicates that the model performed well in early and mid-season (Fig. 5b). In late season, consistent overestimation was observed for all three cultivars and at both sites (Fig. 5b). The comparison of the predictions and the on-site measurements for the other cultivars are shown in Fig. 5c. Overall, the performance of the Auto-ML model ranged between 𝑅𝑀𝑆𝐸 = 1.19 °C and 𝑅𝑀𝑆𝐸 = 2.20 °C, except for ‘Marquette’ whose 𝑅𝑀𝑆𝐸 = 3.20 °C (Fig. 5c). Since the observed LT_50_ of ‘Marquette’ from mid to late season are much higher than the observed LT_50_ of other hybrid cultivars, which is abnormal, we assume the underperformance of the Auto-ML model on ‘Marquette’ might be from repeated measurement errors or sample collection errors. Nevertheless, late season overestimation was also observed in most other cultivars, though it was more severe in some hybrid cultivars and *V. vinifera* cultivars with faster deacclimation response (e.g., ‘Gewurztraminer’ and ‘Lemberger’) (Fig. 5c).

Freezing damage potential was estimated based on the temperature gap between predicted LT_50_ and daily minimum temperature. The freezing damage potential over the dormant season is estimated for the Northeastern and Midwestern U.S and updated daily using gridded weather data. Maximum freezing damage potential and maximum freezing tolerance (minimum predicted LT_50_) in the 2022-2023 dormant season of 16 cultivars are shown in Fig. S3 and Fig. S4, respectively.

## 4. Discussion

In this study, we used AutoGluon, an Auto-ML platform, to assist in the development of a multi-site grapevine freezing tolerance model. Results from method suitability and site-transferability tests indicated the feasibility of the feature extraction and modeling method. The resulting Auto-ML model outperformed two previous models in three test cultivars and accurately predicted the freezing tolerance in the 2022-2023 dormant season. The Auto-ML model was deployed in the major cool climate viticultural regions in the U.S. at new grapevine freezing tolerance website, enabling a large-scale real-time simulation of grapevine freezing tolerance in North America for the first time. The development of such a predictive system is crucial to estimating how cultivated plants can and will adapt to changing climatic conditions. Additionally, freezing tolerance prediction will be advantageous for farmers to determine management strategies and select suitable cultivars under existing and changing climatic conditions (Raza *et al*., 2019; Körner, 2021). It’s also advantageous for the rest of the supply chain and the consumers since appropriate and timely mitigation strategies may reduce fluctuations in produce and product supplies.

Diverse modeling approaches, such as empirical modeling and mechanistic modeling, have been intensively used in the history of plant biology. ML-empowered or DL-empowered data-driven modeling has been gaining favor in recent years for its efficiency and high accuracy, especially in “omics” data that exhibit high complexity (Krantz *et al*., 2021). However, modeling of complex physiological changes in plants using local or regional data often results in models that tend to overfit. Additionally, mechanistic and DL-empowered models can be complicated, reducing the user-friendliness and ultimately adoption (Ellis *et al*., 2020). Detailed discussion of the reasoning for these limitations is included in Note S3. As a novel approach designed to address both issues (tendency of overfitting and easiness of modeling), we collected training data from multiple regions and used Auto-ML to model grapevine LT_50_. The accumulation of grapevine LT_50_ data across North America presented in this study was likely to generate a generalizable model for different climate conditions, and Auto-ML would in turn ease the modeling. As the aim is to generate a site-transferable grapevine LT_50_ prediction model, the modelling engine (AutoGluon) and the features were first evaluated for their suitability to model grapevine LT_50_ and their capacity to generate a stable prediction model with high site-transferability. During the method suitability and site-transferability tests, the alpha model generated using partial data resulted in a similar performance in internal testing (𝑅𝑀𝑆𝐸 = 1.35 °C) and external testing (𝑅𝑀𝑆𝐸 = 1.59 °C) (Fig. S2). These results demonstrated an improvement over predictions produced by the NYUS.1, WAUS.2 and the RNN model during the testing on unseen data (Saxena *et al*., 2022; Kovaleski *et al*., 2023), indicating that the alpha model passed the method suitability test and the site-transferability test. As the internal and external testing data are from different sites, this result also suggests that the AutoGluon method did not overfit the training data and exhibited high site-transferability.

Among the different potential models trained on the entire dataset, ‘WeightedEnsemble_L2’ exhibited high accuracy and low complexity (thus low inference latency) and was chosen as the final model to deploy. Internal testing showed that the performance of the final model (Auto-ML model) varied in different regions (Fig. 3b); however, the regions that show greater 𝑅𝑀𝑆𝐸 have more complex data composition (e.g., a mix of hybrid cultivars and *V. vinifera* cultivars in NY dataset), indicating potential model underperformance in some cultivars. The Auto-ML model in general, outperformed previous models (Fig. 4a). Prediction error distribution showed that NYUS.1 and WAUS.2 lost prediction power when tested in disparate regions. For example, WAUS.2 performs well in WA sub-dataset, however, the predictions in other sub-datasets exhibit frequent overestimation (Fig. 4b). Similarly, NYUS.1 accurately predicts the LT_50_ in NY sub-dataset, yet the prediction of WA and BC sub-datasets are less accurate (Fig. 4b). The underperformance of these two models is more severe in late season as the buds deacclimating and approaching budbreak (Fig. 4b). To compare, the Auto-ML model leverages the multi-region training dataset and generalizes across different regions, which resulted in a more evenly distributed prediction error in different regions (Fig. 4b). This result again indicates that the final model is likely a generalizable model that exhibited high site-transferability.

However, the Auto-ML model also has several limitations. First, the number of parameters in the Auto-ML model is considerably more than the mechanistic models, which might explain its better performance as the same process was formulated with more factors. However, utilizing more parameters might also impair the efficiency of deployment as more computational resources are needed, limiting the potential adoption of this model to users with computational support. Unlike the mechanistic models, in which each prediction is explainable by the parameters, the Auto-ML method utilizes features in a complicated way through stacking and weighting of multiple base models (the black box effect), which impairs the ability to attribute model changes to biological or physiological factors. The black box effect of these models brings limited insight towards understanding grapevine dormant season biology (Azodi *et al*., 2020).

Nevertheless, the impact of each feature can be determined using AutoGluon and SHAP analysis, thus partially unboxing the black box of the model. Although the result differs due to different computational approaches, five features (the Utah chilling accumulation, days in season, EWMA of mean temperature with window size of 12, the NC chilling accumulation, and the GDH with 10 °C as base temperature) were identified for their importance by both algorithms (Fig. 3c and d). These features have much higher statistical importance for the model than the other features, however, the model prediction is not fully explainable using these features only. In fact, the Auto-ML model prediction incorporates complicated interactions of feature groups, and each group contributes to the final prediction in different ways. Detailed analysis and reasoning were included in Note S4 and Fig. S5.

Among these five features, Utah and NC are two chilling models that calculate the accumulation of chilling in the season. Chilling accumulation has been determined for its importance in the physiology of grapevine in the dormant season (Dokoozlian, 1999; Londo & Kovaleski, 2017, 2019; Kovaleski *et al*., 2018). Different chilling models also help quantify the dynamics of cold acclimation and deacclimation in the NYUS.1 and WAUS.2 models (Ferguson *et al*., 2014; Kovaleski *et al*., 2023), and therefore errors from chilling models can hinder predictions. The Auto-ML model utilized individual chilling models in a different manner than the previous prediction models (Fig. 3c and Fig. S6a). The SHAP value of the Utah chilling model exhibits an ‘S’ shape as Utah chilling increases, while NC and CU chilling models exhibit ‘U’ shapes (Fig. S6a). A potential explanation for the importance of both Utah and NC chilling models in Auto-ML is that the Auto-ML model combined the effect of the chilling models and generated a unique algorithm to dynamically describe the effect of chilling accumulation. By simply summing the SHAP values of three chilling models and averaging their normalized feature values, their combined effect reveals a standard ‘U’ shape of grapevine LT_50_ across a dormant season (Fig. S6b). In the early season when the combined chilling unit accumulation is low, it negatively impacts freezing tolerance (positive SHAP). As the combined chilling unit increases, the negative impact gradually decreases, transitions to a positive impact, and remains relatively unchanged after reaching a turning point (Fig. S6b). The turning point (derivative of the loess fitted function 𝑓^′^(𝑆𝐻𝐴𝑃) = 0), occurs at 44% of average normalized feature values, corresponding to 1,245 chilling units in averaged Utah, NC, and CU models (Fig. S6c). This value coincides with the amount of chilling needed to fulfill chilling requirements for most grapevine cultivars (Londo & Johnson, 2014). This result suggests that, although the black box effect of the Auto-ML model impairs its explainability, some biological components of grapevine physiology in the dormant season were captured and utilized by the model.

Another issue of the Auto-ML model is the inconsistency of its performance for different cultivars and different times of the season. Site-transferability tests and model comparison results suggest that the Auto-ML model underperforms for some hybrid cultivars (Fig. S2 and Fig. 4). This underperformance compounds with a prediction that is more error-prone in late season (Fig. 5). Due to the nature of Auto-ML, the mechanism resulting in this lower predictability is not explainable. The only ways to avoid these issues are either training the model with more on-site measurement data or generating more climate descriptors and cultivar features to better characterize weather conditions and individual cultivars in late season.

To our knowledge, our study represents the first attempt at modeling weather-related perennial crop physiology using only an Auto-ML modeling engine. The current model is named NYUS.2 based on the previously proposed model naming system (Kovaleski *et al*., 2023). The method was validated, and the final model revealed high site-transferability and outperformed two previous mechanistic models in all testing regions. Feature importance analysis indicates that the model might have adequately extracted some biological mechanisms during training. The effect of chilling models in the Auto-ML model might indicate the potential identification of chilling fulfillment of grapevine, and the effect one-hot encoded cultivar features might indicate the proper differentiation of cultivar-specific acclimation and deacclimation dynamics (Londo & Johnson, 2014; Kovaleski *et al*., 2018; Londo & Kovaleski, 2019; North *et al*., 2022). Deployment of NYUS.2 in the 2022-2023 dormant season provided daily updated model estimates of grapevine freezing tolerance of 16 cultivars at 2,035 independent locations in 23 states in the U.S. using the weather station database ACIS (https://www.rcc-acis.org/). NYUS.2 and the Shiny-based web application represent the launch of the first large-scale real-time grapevine freezing simulation system with high temporal resolution in North America. In ongoing development, we plan to further expand the deployment of the NYUS.2 model to more cool climate viticultural regions in North America, such as Quebec, Ontario and British Columbia in Canada. Further, then incorporation of more worldwide on-site measurements of freezing tolerance and more climate descriptors would enable further training of the model, thus potentially yielding with a more generalizable model to help understand the biology of grapevine freezing tolerance and quantify the threat of freezing under climate change.

## 5. Data availability

All the original training data and source code for feature extraction, modeling training and model deployment are available at https://github.com/imbaterry11/NYUS.2

## Supporting information

Table S1, Fig. S1-S6 and Note S1-S4 will be used for the link to the file on the preprint site

Data S1-S6 will be used for the link to the file on the preprint site

## 6. Acknowledgements

The authors would like to thank Lynn Mills (WA), Beth Ann Workmaster (WI), Katherine Benedict (NS), Alexander Campbell and Jessee Tinslay (QC), Don Smith and Meredith Persico (PA) and Hanna Martins, Felex Pike, and Bill Wilsey (NY) for their help in LT_50_ data collection. This work was partially supported by the Office of the Vice Chancellor for Research and Graduate Education at the University of Wisconsin–Madison with funding from the Wisconsin Alumni Research Foundation, and through the USDA ARS appropriated project 1910–21220– 006–00D and New York Wine and Grape Foundation. The NY LT_50_ data collection was also supported by Cornell University Federal Capacity Funds Grant Program. The WA LT_50_ data collection was supported by the Washington Wine Industry Foundation. The PA LT_50_ data collection was supported by the USDA National Institute of Food and Agriculture (NIFA) Federal Appropriation under Projects PEN0 4794 (7003432). The NS LT_50_ data collection was supported by a Canadian Agricultural Partnership (CAP) project (ASC-12 Wine Grape Cluster Activity 7), the Canadian Grapevine Certification Network (CGCN) and the Grape Growers’ Association of Nova Scotia (GGANS). The QC LT_50_ data collection was supported by AgriScience program-cluster on behalf of Agriculture and Agri-Food Canada.

## 8. Author contributions

HW and JPL assembled the dataset, conducted the modeling, did the model analysis and designed the website. GDM provided technical and biological guidance of machine learning and feature importance quantification. APK provided the code for WAUS.2 and NYUS.1. APK, MK, TEM, AHW, JLF, AHH, CP, MR, AA, MGN, PW and MC collected LT_50_ data and contributed the sub-datasets from different regions. HW, JPL and GDM wrote a majority of the manuscript with contributions from all co-authors.

