## Supplementary material for "NYUS.2: an Automated Machine Learning Prediction Model for the Large-scale Real-time Simulation of Grapevine Freezing Tolerance in North America": Table S1, Fig. S1-S6 and Note S1-S4 will be used for the link to the file on the preprint site

***New Phytologist* Supporting Information**

Article acceptance date: Click here to enter a date.

**Table S1 Data collection site detail**

**Fig. S1 Correlation analysis of EWMA and REWMA of daily minimum temperature**

**Fig. S2 Result of the method suitability test and the site-transferability test**

**Fig. S3 Maximum potential that freezing damage has occurred in 16 grapevine cultivars in the 2022-2023 dormant season**

**Fig. S4 Maximum freezing tolerance of 16 grapevine cultivars in the 2022-2023 dormant season**

**Fig. S5 Impact of different feature groups on Auto-ML model prediction**

**Fig. S6 Auto-ML model utilization of three chilling models**

**Note S1 Daily temperature descriptors and cumulative temperature descriptors**

**Note S2 Reverse exponential weighted moving average temperatures in the modeling of grapevine freezing tolerance**

**Note S3 Limitations of mechanistic modeling and DL-empowered modeling in plant physiology**

**Note S4 Feature importance investigation based on feature groups**

The following Supporting Information is available for this article:

### Table S1 Data collection site detail

| | Site detail | | | | Cultivar detail | | |
|---|---|---|---|---|---|---|---|
| Location | longitude | latitude | Weather data | Station ID | Cultivar number | Record | Cultivars |
| BC | -119.60 | 49.46 | DACCD[1] | Penticton A | 15 | 2012-11-01 to 2022-04-07 | Cabernet Franc, Cabernet Sauvignon, Chardonnay, Gewurztraminer, Malbec, Merlot, Pinot blanc, Pinot gris, Pinot noir, Riesling, Sauvignon blanc, Syrah, Tempranillo, Viognier, Zinfandel |
| NY | -77.03 | 42.88 | NEWA[2] | Bejo | 32 | 2012-10-31 to 2022-03-07 | Aromella, Cabernet Franc, Cabernet Sauvignon, Cayuga White, Chambourcin, Chancellor, Chardonnay, Chenin blanc, Concord, Corot noir, Gewurztraminer, Gruner Veltliner, La Crescent, Lemberger, Marechal Foch, Marquette, Merlot, Niagara, Noiret, Pinot gris, Pinot noir, Riesling, Sangiovese, Saperavi, Sauvignon blanc, St. Croix, Syrah, Tocai Fruliano, Traminette, Valvin Muscat, Vidal, Vignoles |
| NS | -64.32 | 44.93 | On-site measurement[3] | - | 5 | 2018-10-29 to 2023-02-22 | Chardonnay, Marquette, Pinot noir, Riesling, L'Acadie |
| PA | -77.95 | 40.71 | NEWA | Rock Springs & Lewisburg (FERO vineyards) | 4 | 2018-11-06 to 2022-11-29 | Lemberger, Marquette, Noiret, Riesling |
| WA | -119.74 | 46.25 | Agweather | Prosser. NE & Paterson. E | 12 | 2005-09-28 to 2012-04-17 | Cabernet Franc, Cabernet Sauvignon, Chardonnay, Chenin blanc, Gewurztraminer, Lemberger, Merlot, Pinot gris, Riesling, Sangiovese, Sauvignon blanc, Syrah |
| QC | -74.05 | 45.49 | On-site measurement | - | 13 | 2020-10-20 to 2022-12-13 | Cabernet Franc, Chardonnay, Marquette, Pinot gris, Pinot noir, Riesling, Vidal, Frontenac, Petite Pearl, Frontenac blanc, Frontenac gris, Seyval, St. Pepin |
| MI | -86.36 | 42.08 | Enviroweather | SWMREC | 11 | 2021-11-23 to 2022-04-27 | Cabernet Franc, Cabernet Sauvignon, Concord, Marechal Foch, Marquette, Merlot, Niagara, Pinot gris, Pinot noir, Sauvignon blanc, Traminette |
| TX | -101.82 | 33.65 | ACIS[4] | USW00023042 | 4 | 2021-12-08 to 2022-02-16 | Cabernet Sauvignon, Sangiovese, Tempranillo, Viognier |
| WI | -89.53 | 43.06 | ACIS | USW00014837 | 5 | 2017-11-02 to 2020-04-17 | La Crescent, Marquette, Brianna, Frontenac, Petite Pearl |

[1]Digital Archive of Canadian Climatological Data (DACCD)

[2]Network for Environment and Weather Applications (NEWA)

[3]Weather data was obtained from on-site weather stations

[4]Applied Climate Information System (ACIS)

**Fig. S1** Correlation analysis of EWMA and REWMA of daily minimum temperature. All the EWMA and REWMA of daily minimum temperatures in the entire dataset was used for the correlation analysis.

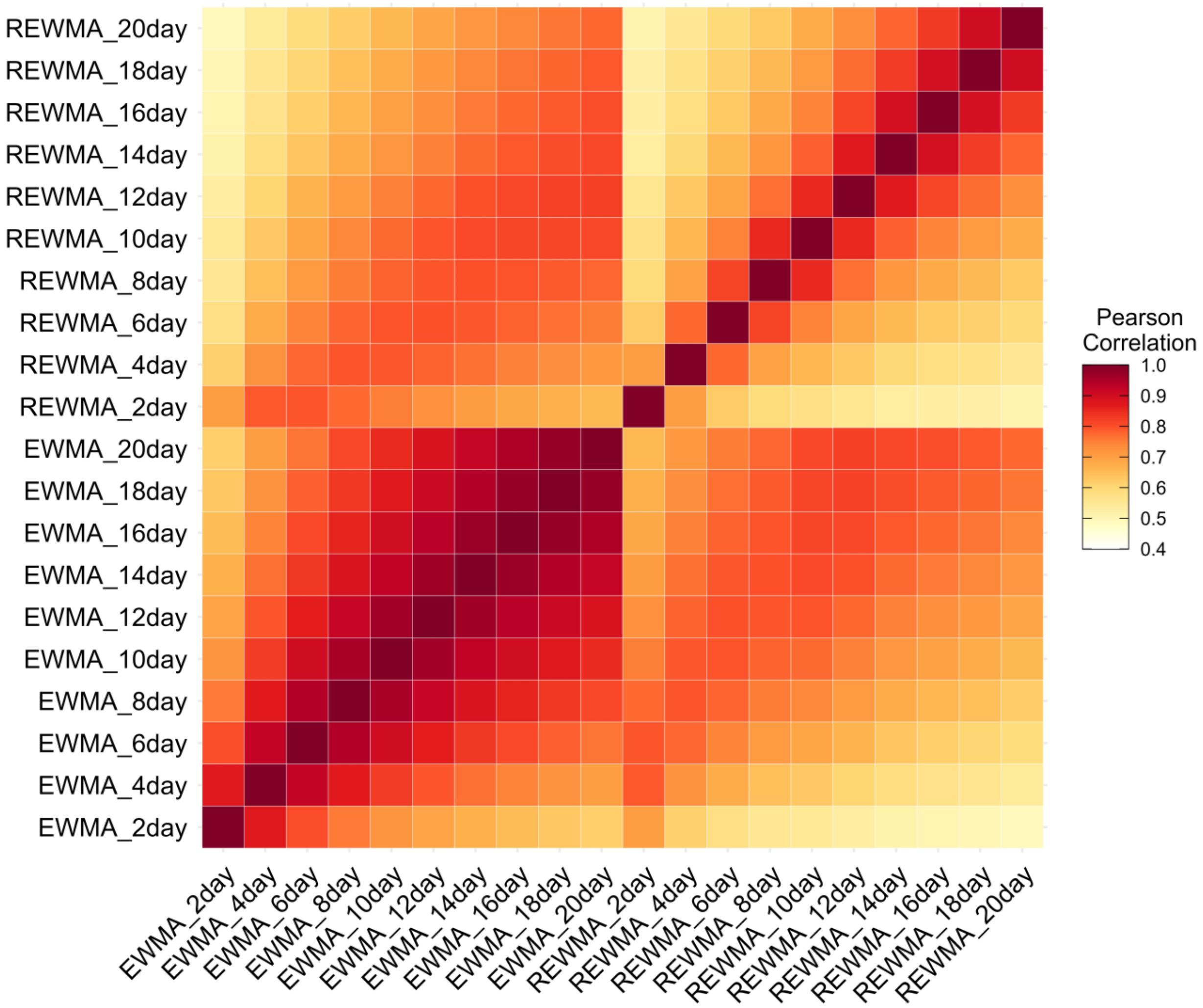

**Fig. S2** Result of the method suitability test and the site-transferability test. (a) The performance (RMSE) of all the models generated from training on internal testing data ($n = 970$). Inference latency refers the relative time it takes for a machine learning model to process and analyze data and produce an output. (b) The performance of the alpha grapevine $LT_{50}$ prediction model on external testing data ($n = 459$) by individual sub-datasets and cultivars.

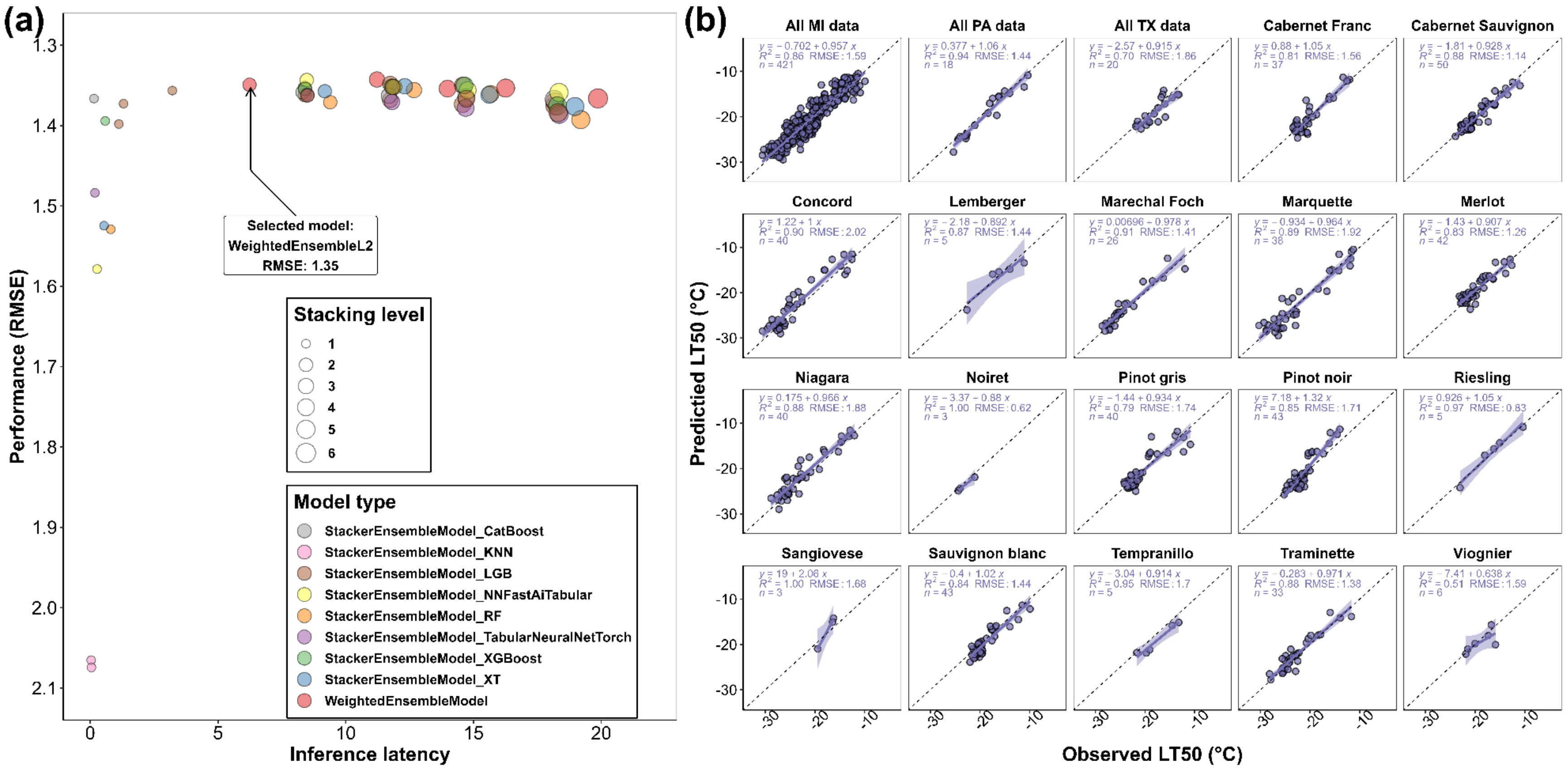

**Fig. S3** Maximum potential that freezing damage has occurred in 16 grapevine cultivars in the 2022-2023 dormant season. The potential that a freezing damage has occurred (0 to 100%), is estimated through a symmetric sigmoid function assuming that 10% and 90% of potential that a freezing damage has occurred when the ambient temperature is 2 °C above and below the predicted $LT_{50}$, respectively

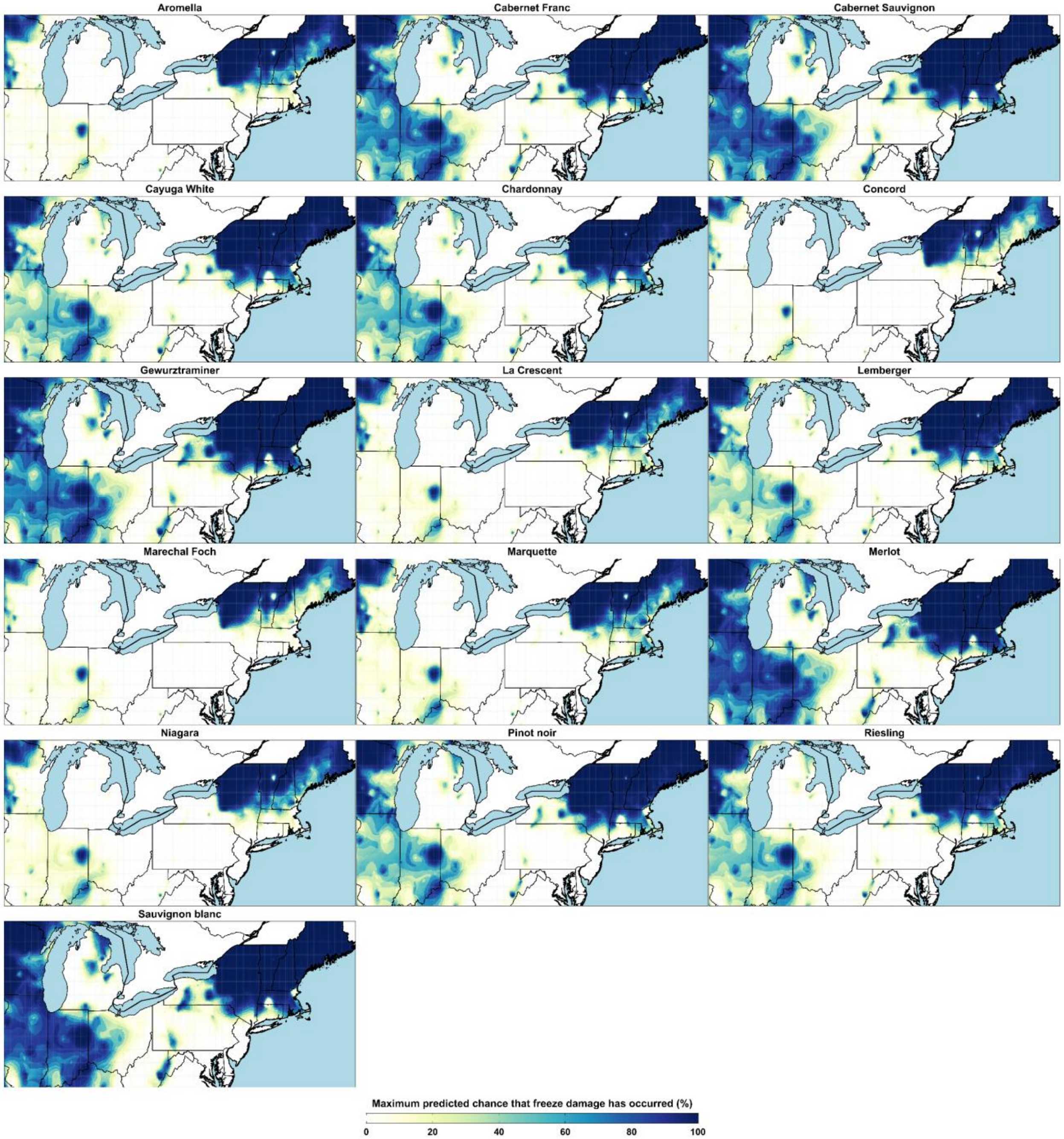

**Fig. S4** Maximum freezing tolerance of 16 grapevine cultivars in the 2022-2023 dormant season. The freezing tolerance is expressed as $LT_{50}$ (°C).

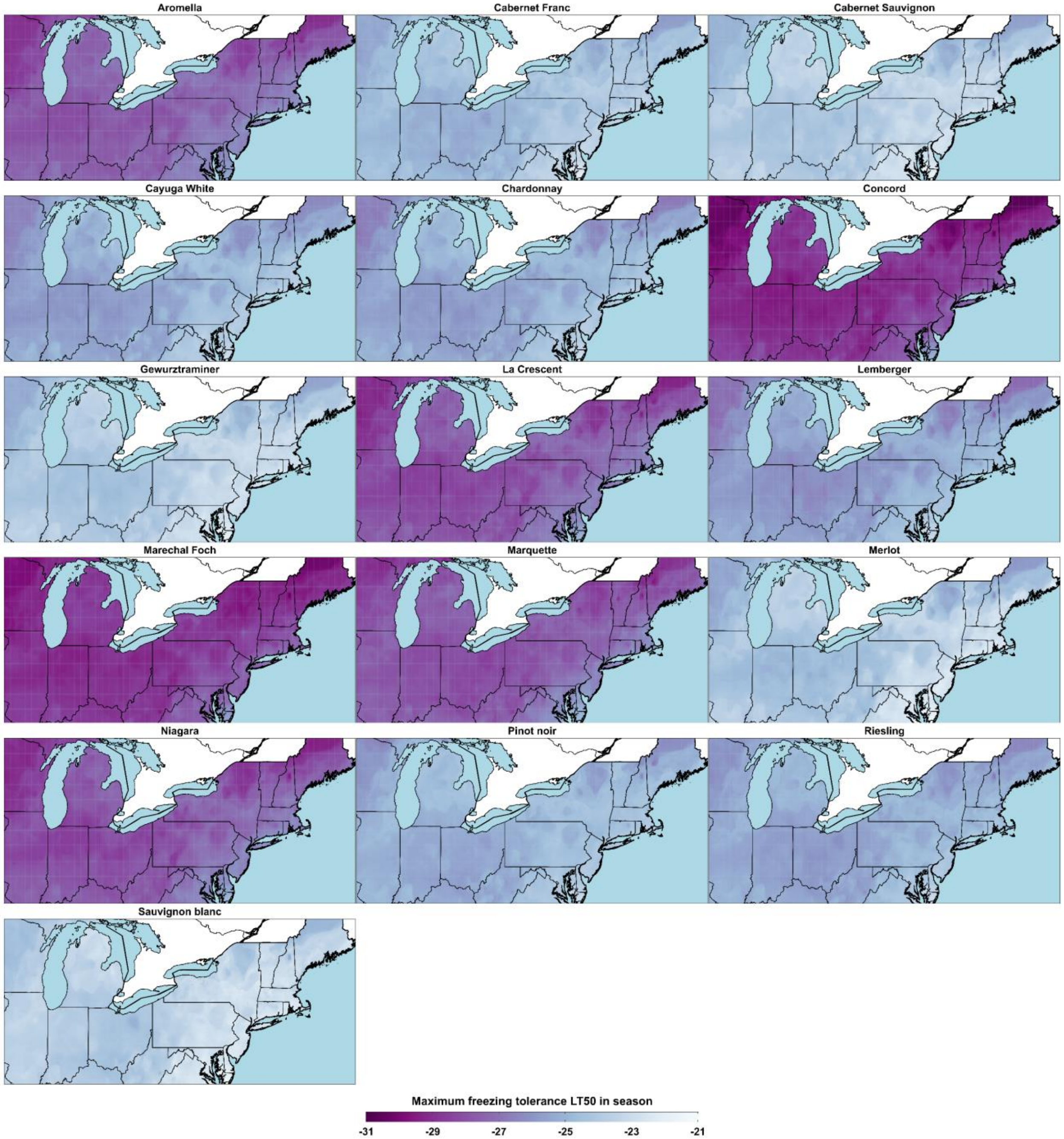

**Fig. S5** Impact of different feature groups on Auto-ML model prediction. (a) Impact of top features on model prediction. The top features include the five shared features among the top 15 important features quantified by AutoGluon and SHAP value. (b) Impact of EWMA and REWMA temperatures on model prediction. EWMA and REWMA temperatures include 60 features computed with different moving average methods using daily temperatures. (c) Impact of cultivar features on model prediction. Cultivar features include 45 one-hot encoded cultivars. Internal testing data ($n = 1{,}016$) was used for the model prediction comparison. The plots are showing the prediction of the $LT_{50}$ in Geneva, NY in the 2022-2023 dormant season.

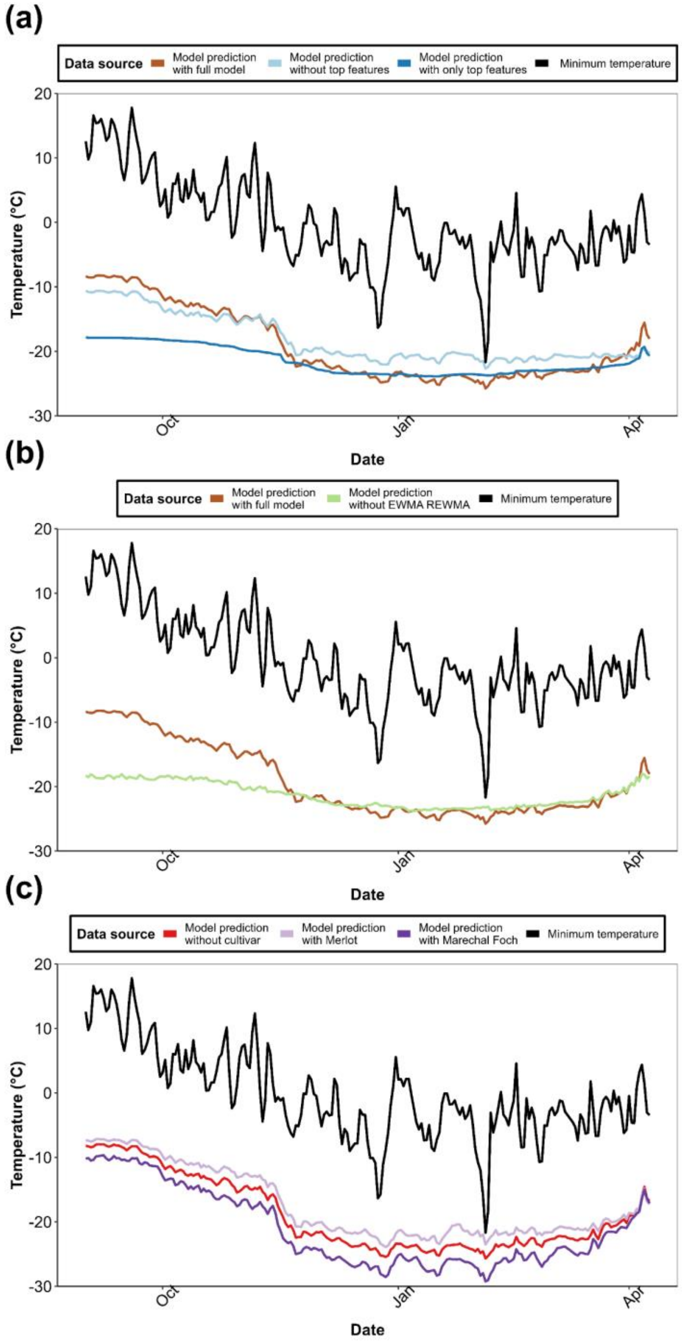

**Fig. S6** Auto-ML model utilization of three chilling models. (a) The change of SHAP value with the increase of normalized feature values in three chilling models. Normalized feature value is calculated with $x_{normalized} = (x - x_{\min})/(x_{\max} - x_{\min})$ and ranged from 0 to 1. (b) The change of combined SHAP value with the increase of combined normalized feature value. Combined SHAP value is calculated by summing the SHAP value of the three chilling models. Combined normalized feature value is calculated by average the normalized feature values of the three chilling models. The trendline is fitted using 'loess' with $\alpha = 0.7$. (c) Derivative of the fitted trendline over combined normalized feature value.

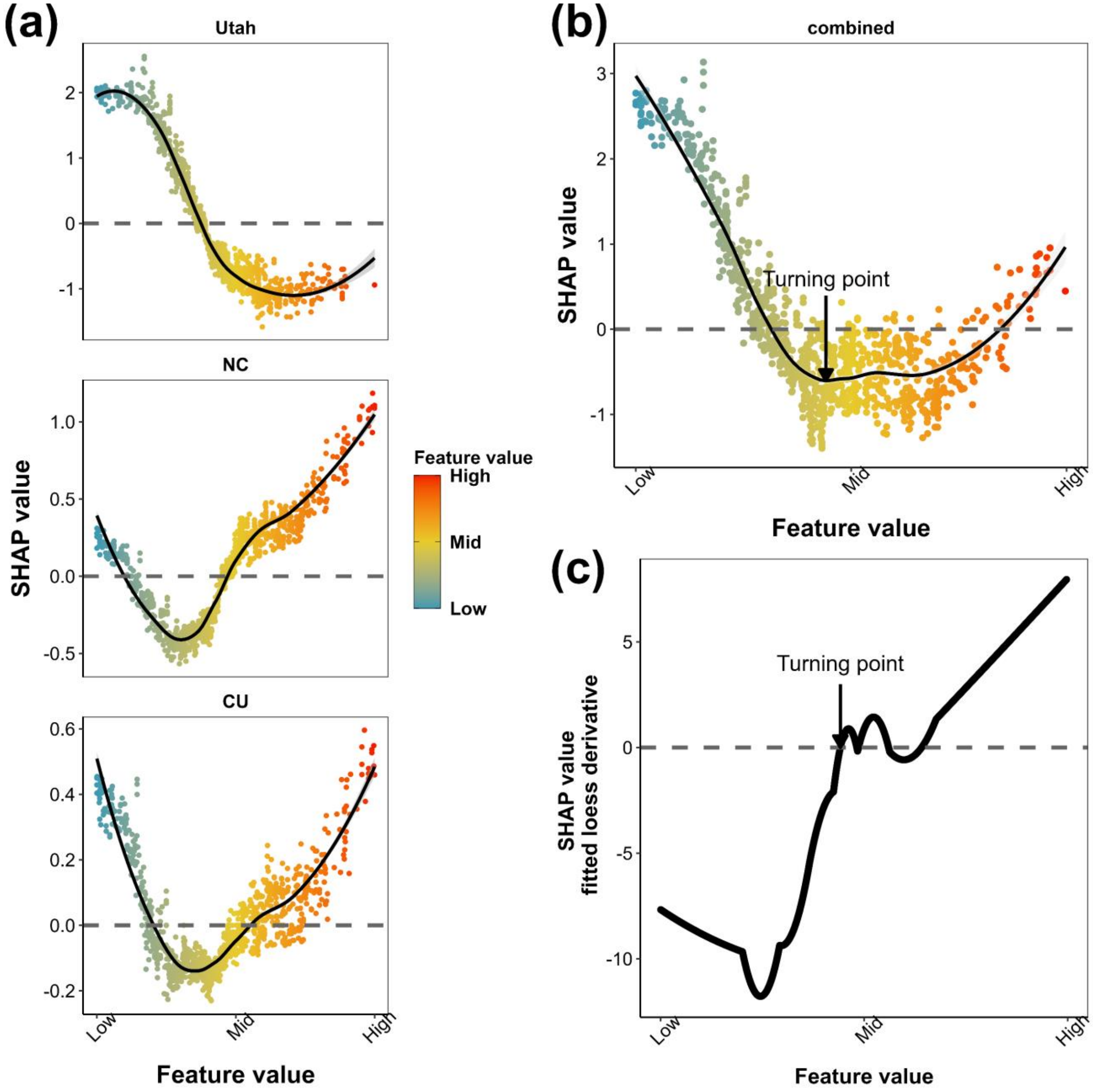

**Note S1** Daily temperature descriptors and cumulative temperature descriptors

Daily temperature descriptors for the modeling of grapevine freezing tolerance in this study include daily maximum temperature, daily minimum temperature, daily mean temperature and within-day temperature range. Cumulative temperature descriptors include chilling units as estimated from different models and growing degree hours (GDH) with different base temperatures to account for the accumulation of chill and heat throughout the season. Chilling units were estimated with three chilling models, the Chilling Hours model (CU), the Utah model (Utah), and the North Carolina model (NC), using September 1$^{st}$ in each year as the start of dormant season (Weinberger, 1950; Shaltout & Unrath, 1983; Linvill, 1990; Camargo-Alvarez *et al.*, 2020). GDHs were computed with the base temperature at 10 °C, 7 °C, 4 °C and 0 °C, starting from January 1$^{st}$ in each year, as a standard approach for the modeling of phenology in perennial plants (Diekmann, 1996; Fu *et al.*, 2014; Zapata *et al.*, 2017). Cumulative temperature descriptors were computed using R packages 'chillR' and 'fruclimadapt' (Luedeling & Fernandez, 2022; Miranda, 2023).

**Note S2** Reverse exponential weighted moving average temperatures in the modeling of grapevine freezing tolerance

REWMA temperature is developed in this study to better correlate the physiological response of temperature with grapevine freezing tolerance by formulizing a theory of 'cold priming' and the 'cold shock effect' through modified EWMA. The cold priming theory is founded on the observations that plants that have been cold-primed and then allowed to recover at warmer temperatures are able to gain freezing tolerance more quickly to cold stress than the plants that have not undergone the initial cold exposure (Schwachtje *et al.*, 2019; Leuendorf *et al.*, 2020; Sharma *et al.*, 2022; Liu *et al.*, 2022; Babajamali *et al.*, 2022). Cold priming was first reported in annual plants such as *Arabidopsis thaliana* and *Brachipodium distachyon* (Zuther *et al.*, 2019; Mayer & Charron, 2021). Factoring cold priming in the modeling of *Citrus sinensis* freezing tolerance also resulted in a high accuracy dynamic prediction model (Kimura *et al.*, 2021). Assuming that cold priming also exists in grapevine, the freezing tolerance in grapevine on a date might not only be modulated by the continuous exposure to a temperature window that are closed to the date (which is addressed using EWMA) but also be impacted by earlier temperature deviants, which might have induced cold priming. Moreover, in grapevine dormant season physiology, earlier condition might sometimes have more impact than current condition, especially during the early stages of cold acclimation when buds are endodormant. A cold shock in early season would significantly enhance the freezing tolerance of endodormant grapevine, and this enhancement would not fade even under a following period of higher temperature. The phenomena is partially explained by the cold acclimation and deacclimation dynamics during accumulation of chilling units (Kovaleski *et al.*, 2018, 2023). However, since we aim to develop a temperature-based prediction model rather than a biology-based prediction model, we should include sufficient thermal features for Auto-ML to factor these theories so that it might identify the underlying biology. To show the potential effect of early sudden temperature abnormality on current freezing tolerance, we exponentially added more weight to earlier temperatures when computing moving average. Compared to the computation of EWMA, the assignment of weight is reversed, thus this method is named REWMA (reverse EWMA).

**Note S3** Limitations of mechanistic modeling and DL-empowered modeling in plant physiology

The leading issues with modeling plant physiology are usually the availability of training data and the ease of modeling itself. The WAUS.2 model is a product of mechanistic modeling that mathematically realizes an understanding of grapevine dormant season biology (Ferguson *et al.*, 2014). The NYUS.1 model uses a combination of mechanistic and empirical modeling through the formulation of empirically determined concepts of grapevine dormant season physiology (Kovaleski *et al.*, 2023). Smaller training datasets are sufficient for training and generating such models. However, the models trained using such datasets tend to overfit niche conditions. In contrast, the RNN grapevine $LT_{50}$ prediction model is a product of DL, in which unstructured weather data in the whole season are used for training (Saxena *et al.*, 2022). However, generally, tens of thousands of training data are required to generate a stably functional model for plant physiology (Soltis *et al.*, 2020; van Dijk *et al.*, 2021; Gall *et al.*, 2022). Thus, a potential challenge of this model is the potential underperformance due to the relatively low quantity of training data ($n < 8{,}000$). The RNN model cannot outperform WAUS.2 in some cultivars under Washington state weather conditions (Saxena *et al.*, 2022).

Mechanistic models and DL-empowered models are complicated to build, but the underlying reasons are different. Current constraints on traditional mechanistic modeling methods include the need for extensive and manual inputs and training requirements for both developers and end-users. For developers, model construction involves a thorough understanding of the biological systems for parameter selection and combination, which often results in complicated models that require specialized expertise to interpret (Cartwright *et al.*, 2016). For users, these issues can make mechanistic models less user-friendly and less likely to be adopted (Ellis *et al.*, 2020). Regarding DL-empowered modeling, although the modeling itself involves less biological reasoning, the model selection, the design of the neural networks backbone, and the hyperparameter selection require intensive manual tuning by computer scientists and specialized computational hardware (e.g., GPUs), thus reducing adoption in the field of plant physiology (Janiesch *et al.*, 2021).

**Note S4** Feature importance investigation based on feature groups

As we noticed that several most important features from feature importance quantification exhibited much higher importance scores than the other features, we precisely analyzed if the model only relies on these top features for prediction. The impact of two other groups of features, EWMA and REWMA temperature features and cultivar features, were also analyzed. Internal testing data $n = 1{,}016$ were used for model prediction and performance comparisons between the full model and the models without certain feature groups.

The top features selected for the analysis are the five shared features from top 15 most important feature lists generated by AutoGluon and SHAP value. These features are the chilling accumulation estimation of Utah model, days in season, the EWMA of mean temperature with window size of 12, the chilling accumulation estimation of NC model, and the GDH with 10 °C as base temperature. By excluding these top features, the $RMSE$ of the model prediction on internal testing data increased from 1.36 °C (full model prediction) to 3.01 °C. By including only these top features, the $RMSE$ of the model prediction on internal testing data increased to 4.12 °C. An example of model prediction difference between the models without/with only the top features and the full model is shown in Fig. S5a. Compared to the prediction from the full model, the prediction without the top features followed the same trend during cold acclimation in early season and the maintenance of maximum freezing tolerance in mid season but did not reveal the response of deacclimation in late season (Fig. S5a). Compared to the prediction from the full model, the prediction with only these top features neither responded to daily temperature variations nor exhibited accuracy in early season (Fig. S5a). These results indicate that, the top features are essential for the prediction of LT50 during deacclimation, however, the model does not only rely on these features for prediction.

The analysis of EWMA and REWMA temperature features incorporated the comparison between the performance of the model without 60 EWMA and REWMA temperature features and the full model. By excluding the EWMA and REWMA features, the $RMSE$ of the model prediction on internal testing data increased to 4.12 °C. The model without the EWMA and REWMA features lost prediction accuracy during cold acclimation in early season and did not reveal proper response to daily temperature changes (Fig. S5b). The analysis of cultivar features incorporated the comparison between the performance of the model without 45 one-hot encoded cultivar features and the full model with proper cultivar features. The impact of using proper cultivar feature on model prediction is shown in Fig. S5c, using the cultivars ‘Marechal Foch’ and ‘Merlot’ as examples. By excluding all the cultivar features, the model could not generate cultivar-specific prediction, and the predicted $LT_{50}$s are most likely intermediate $LT_{50}$s generalized among all the cultivars, thus might be used as a backbone to generate cultivar-specific prediction (Fig. S5c). To compare, when adding proper cultivar features to the model, the predicted $LT_{50}$s are calibrated to reveal cultivar-specific responses to temperature (Fig. S5c). For example, ‘Marechal Foch’ tends to acclimate faster in early season, maintain lower $LT_{50}$ in mid season and deacclimates faster in late season, as compared to ‘Merlot’ (Fig. S5c). These results align with previous findings regarding the differentiation of acclimation and deacclimation dynamics in different grapevine cultivars, indicating that the cultivar-specific responses in freezing tolerance are addressed by the Auto-ML model with the cultivar features (Kovaleski *et al.*, 2018; North *et al.*, 2022).
